# Prediction of cephalosporin and carbapenem susceptibility in multi-drug resistant Gram-negative bacteria using liquid chromatography-tandem mass spectrometry

**DOI:** 10.1101/138594

**Authors:** Yuiko Takebayashi, Wan Ahmad Kamil Wan Nur Ismah, Jacqueline Findlay, Kate J. Heesom, Jay Zhang, O. Martin Williams, Alasdair P. MacGowan, Matthew B. Avison

**Affiliations:** School of Cellular & Molecular Medicine, University of Bristol, Bristol. UK; Faculty of Biotechnology & Biomolecular Sciences, Universiti Putra Malaysia, Selangor Darul Ehsan, Malaysia; Bristol Proteomics Facility, University of Bristol, UK; Department of Microbiology, Southmead Hospital, Bristol, UK.; Department of Microbiology and Infectious Diseases, Bristol Royal Infirmary, Bristol, UK

**Author notes:** Correspondence: Matthew B. Avison, School of Cellular & Molecular Medicine, Biomedical Sciences Building, University Walk, Bristol. BS81TD. UK.

## Abstract

In vitro antibacterial susceptibility testing informs clinical decision making concerning antibacterial therapeutics. Predicting, in a timely manner, which bacterial infection will respond to treatment by a given antibacterial drug reduces morbidity, mortality, and healthcare costs. It also allows prudent antibacterial use, because clinicians can focus on the least broad-spectrum agent suitable for each patient. Existing susceptibly testing methodologies rely on growth of bacteria in the presence of an antibacterial drug. There is significant interest in the possibility of predicting antibacterial drug susceptibility directly though the analysis of bacterial DNA or protein, because this may lead to more rapid susceptibility testing directly from clinical samples. Here we report a robust and tractable methodology that allows measurement of the abundance of key proteins responsible for antibacterial drug resistance within samples of 1 µg of total bacterial protein. The method allowed correct prediction of β-lactam susceptibility in clinical isolates from four key bacterial species and added considerable value over and above the information generated by whole genome sequencing, allowing for gene expression, not just gene presence to be considered, which is key when considering the complex interplays of multiple mechanisms of resistance.

## INTRODUCTION

Antibacterial drug resistance (ABR) is one of the most serious problems facing mankind [1]. In the developed world, where ABR is currently most significant in the context of healthcare associated infections, it is particularly relevant for bacteria from the “ESKAPE” group (*Enterococcus spp*., *Staphylococcus aureus, Klebsiella pneumoniae, Acinetobacter baumannii, Pseudomonas aeruginosa* and *Enterobacter spp*.) plus *Escherichia coli* [2]. Bacteria that are resistant to all currently available antibacterials exist and this conjures up a nightmare future of untreatable infections. However, one of the most pressing dangers of ABR is that patients with serious infections may not be given appropriate therapy soon enough to avert their deterioration and subsequent death [3]. Because of the wide range of different ABR mechanisms carried by bacteria, empiric therapy is moving towards the use of last resort drugs to which resistance is less likely [4]. However, there is no extant antibacterial to which resistance is never seen, and increasing reliance on last resort drugs inevitably applies selective pressure that drives the evolution of their demise [5]. Therefore, tools to help clinicians rapidly switch to targeted therapy will save human lives and help retain last resort drugs for future use.

One way to dramatically reduce the time it takes to provide key information relevant to antibacterial drug therapeutic choice is to identify bacteria, and perhaps even ABR mechanisms in clinical samples without the need for prior culture. PCR can be used to identify certain mobile ABR genes, and PCR based 16S metagenomic sequencing or equivalent can be used to identify the species of bacterium present in a clinical sample. A wider variety of ABR genes might be identified in one assay using microarray hybridisation technology, but whole genome sequencing (WGS) is touted as being a more comprehensive answer to this question [6–11]. In recent years, the speed, accuracy, cost, and the amount of DNA required for WGS have all shifted by orders of magnitude in favour of routine WGS from clinical samples being a real possibility in the medium term and some major successes have been recorded, particularly where bacterial density is high, e.g. urinary tract infection [12]. Importantly, WGS potentially allows the complex interplays between mobile ABR genes and background mutations to be integrated in a prediction of ABR, something that is particularly necessary in Gram-negative bacteria, where ABR phenotype is frequently multi-factorial [13]. However, this spotlights our lack of understanding of the way genotype relates to ABR phenotype. Without a detailed understanding of which mutations influence, and which do not influence ABR, we cannot hope to accurately predict ABR from WGS in all cases, and the need for more research in this area was highlighted in a recent EUCAST sub-committee report [14].

One of the main information weaknesses of using WGS is a lack of understanding about how genotype affects gene expression levels. And for ABR, protein abundance can have a profound effect. For example, In the presence of weak carbapenem hydrolysing β-lactamases, such as CTX-M and CMY, mutations that reduce the rate of carbapenem entry can help confer carbapenem resistance. Such mutations reduce the abundance of the outer membrane porin through which the carbapenem enters, and/or increase the abundance of efflux pump(s) involved in removing the carbapenem from the cell [15]. We have recently shown, for example, that loss of function mutations in the repressor *ramR* in *K. pneumoniae* enhances efflux pump production and reduces porin production, and this reduces carbapenem susceptibly in CTX-M or CMY producers [16]. Potentially *ramR* sequence nucleotide polymorphisms (SNPs) might be identified in WGS data, but then a decision must be made as to whether the SNPs affect protein function or not.

An alternative approach is to measure protein abundance using proteomics methodologies. This has the potential to give both identification and abundance data that might resolve many of the uncertainties surrounding the use of WGS. This has recently been reviewed, and, whilst there have been some successes identifying some ABR proteins in some cases, a methodology to allow whole proteome analysis that can accurately quantify ABR proteins, which can be used to accurately predict antimicrobial susceptibility is yet to be demonstrated [17]. We have been using so called “shotgun” proteomics via a nano-liquid chromatography, Orbitrap tandem mass spectrometry approach to characterise proteomic responses to mutations in regulators that affect ABR [16,18]. Accordingly, without significant adaptation of the methodology, in the work reported here, we attempted to quantify and identify ABR proteins directly from clinical bacterial isolates grown in culture, and use this information to predict antimicrobial susceptibility across a range of species. We have demonstrated that the approach is tractable, and that it reveals novel biology for future study.

## MATERIALS AND METHODS

### Bacterial strains and antibiotic susceptibility testing

Transformants were made using the following strains: *K. pneumoniae* NCTC5055, *E. coli* 17 (a urinary tract isolate, and gift from Dr Mandy Wootton, Public Health Wales), *P. aeruginosa* PA01, *A. baumannii* CIP 70.10. Clinical isolates were forty human bloodstream *K. pneumoniae* isolates, four *E. coli*, five *A. baumannii* and 5 *P. aeruginosa* (collected as part of the SENTRY antimicrobial surveillance programme and a gift from Prof Tim Walsh, Cardiff University). Disc susceptibility testing was performed according to CLSI methodology [19] and interpreted using CLSI performance standards [20].

### Cloning genes, transformation and complementation studies of K. pneumoniae Ecl8

All recombinant plasmids, where β-lactamase genes had been ligated into Enterobacteriaceae-specific vector pSU18 have recently been described [16]. For transformation into non-Enterobacteriaceae, genes were subcloned into the vector pUBYT, being the plasmid pYMAb2 [21] which we modified to remove the OXA promotor region (located upstream of the multiple cloning site) by PCR amplification using the primers listed in Table 1, followed by digestion with XbaI and ligation to produce a circular product. subcloning into pUBYT used the same restriction enzymes as used to originally clone the genes into pSU18 [16]. Inserts were confirmed by PCR and sequencing using the primers listed in Table 1. Recombinant plasmids were used to transform bacteria to chloramphenicol (30 mg/L) – for pSU18 recombinants – or kanamycin (50 mg/L) – for pUBYT recombinants – resistance using electroporation as standard for laboratory-strain *E. coli*, except that for production of competent cells, *A. baumannii* cells were washed using 15% v/v glycerol in water rather than 10% v/v used for the other species.

**Table 1.**
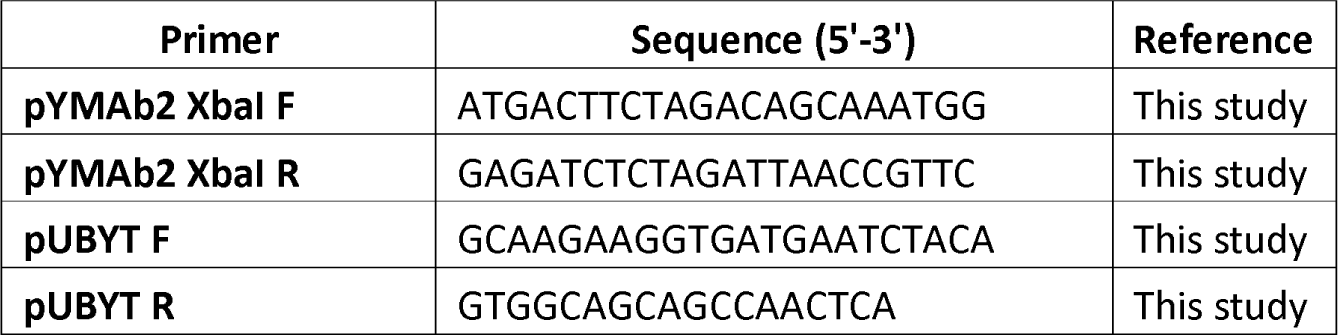
Primers used in this study.

### Quantitative analysis of whole cell proteomes via Orbitrap LC-MS/MS

Each clinical isolate was cultured in 50 ml Cation Adjusted Nutrient Broth (Sigma) with appropriate antibiotic selection. Cultures were incubated with shaking (160 rpm) at 37 C until OD_600_ reached 0.6-0.8. Cells in cultures were pelleted by centrifugation (10 min, 4,000 × *g*, 4°C) and resuspended in 20 mL of 30 mM Tris-HCl, pH 8 and broken by sonication using a cycle of 1 sec on, 1 sec off for 3 min at amplitude of 63% using a Sonics Vibracell VC-505TM (Sonics and Materials Inc., Newton, Connecticut, USA). The sonicated samples were centrifuged at 8,000 rpm (Sorval RC5B PLUS using an SS-34 rotor) for 15 min at 4°C to pellet intact cells and large cell debris and 1 µg of total protein from the supernatant were separated by SDS-PAGE using 11% acrylamide, 0.5% bis-acrylamide (Biorad) gels and a Biorad Min-Protein Tetracell chamber model 3000X1. Gels were run at 200 V until the dye front had moved approximately 1 cm into the separating gel. Proteins in gels were stained with Instant Blue (Expedeon) for 5 min and de-stained in water. The one centimetre of gel lane containing each sample was cut out and proteins subjected to in-gel tryptic digestion using a DigestPro automated digestion unit (Intavis Ltd).

The resulting peptides were fractionated using an Ultimate 3000 nanoHPLC system in line with an LTQ-Orbitrap Velos mass spectrometer (Thermo Scientific) [18]. The raw data files were processed and quantified using Proteome Discoverer software v1.4 (ThermoScientific) and searched against the UniProt *K. pneumoniae* strain ATCC 700721 / MGH 78578 database (5126 protein entries; UniProt accession 272620), the *P. aeruginosa* PA01 database (5563 proteins; UniProt accession UP000002438), the *A. baumannii* ATCC 17978 database (3783 proteins; UniProt accession UP0006737) or the *E. coli* MG1655 database (4307 proteins; UniProt accession UP000000625), in each case, the strain-specific proteome database was augmented by addition of a mobile resistance determinant database (24694 proteins), which was generated by searching UniProt with “IncA”, “IncB” etc to “Inc-Z” as the search term and downloading each list of results. The database file is provided as supplementary data. Proteomic searches against the databases was performed using the SEQUEST (Ver. 28 Rev. 13) algorithm. Protein Area measurements were calculated from peptide peak areas using the “Top 3” method [22] and were then used to calculate the relative abundance of each protein. Proteins with fewer than three peptide hits were excluded from the analysis. Proteomic analysis was performed once for each clinical isolate. For each sample, raw protein abundance for each protein was divided by the average abundance of the 50S and 30S ribosomal proteins also found in that sample to normalise for sample to sample loading variability.

### Whole genome sequencing and data analysis

Genomes were sequenced by MicrobesNG (Birmingham, UK) on a HiSeq 2500 instrument (Illumina, San Diego, CA, USA). Reads were trimmed using Trimmomatic [23] and assembled into contigs using SPAdes 3.10.1 (http://cab.spbu.ru/software/spades/). STs, the presence of resistance genes, and plasmid replicon types were determined using MLST 1.8, ResFinder 2.1, [24] and PlasmidFinder [25] on the Center for Genomic Research platform (https://cge.cbs.dtu.dk/services/).

## RESULTS & DISCUSSION

### Establishing predictive rules for β-lactamase mediated β-lactam resistance

Our initial aim was to generate transformants of the four test species (*K. pneumoniae, E. coli, P. aeruginosa* and *A. baumannii*), each carrying one of seven clinically important β-lactamase genes, each cloned downstream of a natural promoter, representative of promoters found in clinical isolates, and present on a low copy number cloning vector [16]. Once the transformants had been confirmed, we determined the β-lactam susceptibility profile of each (Table 2). These data were used to define rules to predict which β-lactam resistance phenotypes are conferred by carriage of each β-lactamase in each species. We defined these rules in the knowledge that factors affecting β-lactamase abundance and background cell permeability are likely to affect the applicability of these rules, but with the desire to make the rules more flexible by integrating such additional information.

**Table 2.** Antimicrobial susceptibility testing for β-lactamase producing transformants.

| Strain | FOX | CXM | CRO | CTX | CAZ | FEP | ATM | IPM | MEM | DOR | ETP |
| --- | --- | --- | --- | --- | --- | --- | --- | --- | --- | --- | --- |
| K. pneumoniae pSU18 | 23 | 24 | 31 | 33 | 31 | 34 | 34 | 29 | 30 | 28 | 33 |
| K. pneumoniae pSU18 KPC-3 | 18 | 6 | 18 | 19 | 10 | 20 | 6 | 17 | 19 | 16 | 16 |
| K. pneumoniae pSU18 VIM-1 | 9 | 6 | 18 | 13 | 6 | 18 | 31 | 16 | 18 | 14 | 19 |
| K. pneumoniae pSU18 IMP-1 | 6 | 6 | 16 | 16 | 6 | 19 | 33 | 17 | 16 | 15 | 19 |
| K. pneumoniae pSU18 OXA-48 | 23 | 21 | 27 | 30 | 28 | 30 | 31 | 20 | 20 | 18 | 15 |
| K. pneumoniae pSU18 CTX-M | 19 | 6 | 6 | 9 | 14 | 19 | 12 | 24 | 29 | 25 | 27 |
| K. pneumoniae pSU18 CMY-2 | 21 | 6 | 16 | 17 | 17 | 31 | 21 | 29 | 29 | 25 | 29 |
| K. pneumoniae pSU18 NDM-1 | 6 | 6 | 6 | 6 | 6 | 16 | 30 | 13 | 12 | 10 | 12 |
| E. coli pSU18 | 31 | 26 | 33 | 35 | 32 | 38 | 36 | 30 | 34 | 33 | 31 |
| E. coli pSU18 KPC-3 | 28 | 6 | 15 | 24 | 18 | 24 | 16 | 19 | 20 | 20 | 18 |
| E. coli pSU18 VIM-1 | 17 | 6 | 16 | 13 | 10 | 23 | 37 | 22 | 25 | 18 | 37 |
| E. coli pSU18 IMP-1 | 6 | 6 | 12 | 12 | 6 | 17 | 31 | 19 | 17 | 15 | 16 |
| E. coli pSU18 OXA-48 | 28 | 25 | 34 | 32 | 30 | 35 | 29 | 27 | 27 | 28 | 27 |
| E. coli pSU18 CTX-M | 28 | 6 | 6 | 6 | 21 | 19 | 18 | 30 | 31 | 30 | 30 |
| E. coli pSU18 CMY-2 | 19 | 6 | 18 | 18 | 17 | 32 | 21 | 29 | 34 | 30 | 32 |
| E. coli pSU18 NDM-1 | 6 | 6 | 6 | 6 | 6 | 9 | 32 | 11 | 11 | 10 | 10 |
| P. aeruginosa pUBYT | 6 | 6 | 30 | 30 | 37 | 34 | 36 | 32 | 37 | 31 | 24 |
| P. aeruginosa pUBYT KPC-3 | 12 | 6 | 6 | 6 | 6 | 6 | 6 | 13 | 6 | 6 | 6 |
| P. aeruginosa pUBYT VIM-1 | 6 | 6 | 13 | 10 | 14 | 9 | 34 | 12 | 18 | 8 | 13 |
| P. aeruginosa pUBYT IMP-1 | 6 | 6 | 6 | 6 | 10 | 7 | 39 | 11 | 6 | 6 | 6 |
| P. aeruginosa pUBYT OXA-48 | 6 | 6 | 31 | 31 | 39 | 35 | 39 | 15 | 17 | 9 | 9 |
| P. aeruginosa pUBYT CTX-M | 6 | 6 | 6 | 6 | 22 | 12 | 19 | 31 | 26 | 23 | 16 |
| P. aeruginosa pUBYT CMY-2 | 6 | 6 | 8 | 11 | 20 | 29 | 23 | 30 | 26 | 25 | 15 |
| P. aeruginosa pUBYT NDM-1 | 6 | 6 | 6 | 6 | 6 | 6 | 34 | 6 | 6 | 6 | 6 |
| A. baumannii pUBYT | 14 | 22 | 30 | 31 | 35 | 37 | 24 | 44 | 42 | 41 | 26 |
| A. baumannii pUBYT KPC-3 | 11 | 6 | 6 | 6 | 6 | 6 | 6 | 12 | 10 | 9 | 6 |
| A. baumannii pUBYT VIM-1 | 6 | 6 | 6 | 6 | 6 | 6 | 21 | 11 | 13 | 6 | 12 |
| A. baumannii pUBYT IMP-1 | 6 | 6 | 6 | 6 | 6 | 6 | 20 | 10 | 10 | 6 | 6 |
| A. baumannii pUBYT OXA-48 | 13 | 14 | 19 | 18 | 27 | 25 | 19 | 16 | 13 | 14 | 8 |
| A. baumannii pUBYT CTX-M | 12 | 6 | 6 | 6 | 17 | 11 | 6 | 34 | 31 | 30 | 21 |
| A. baumannii pUBYT CMY-2 | 6 | 6 | 6 | 6 | 6 | 23 | 17 | 30 | 30 | 29 | 18 |

Zone diameters are reported in mm, following standard CLSI methodology. Shading: Blue,
susceptible, Green, non-susceptible (intermediate resistant), Yellow, resistant. Grey –
assumed to be intrinsically resistant, so no breakpoint defined.

### Predicting β-lactam susceptibility in K. pneumoniae clinical isolates from LC-MS/MS

Our predictive rules (Table 2) were first used to predict β-lactam susceptibility in 40 *K. pneumoniae* clinical isolates grown in broth culture as shown in Error! Reference source not found̤ The primary data was from LC-MS/MS, confirming the presence/absence of various key β-lactamases. To make this realistic in terms of a potential diagnostic methodology, each sample was analysed by LC-MS/MS only once. WGS was used to validate the LC-MS/MS results, in terms of mobile ABR gene presence. A few issues were noted: SHV is intrinsic to *K. pneumoniae* [26] and all isolates were positive based on WGS, as expected, but few were positive in the LC-MS/MS. However, it is well known that basal transcription from the chromosomal bla_SHV_ promoter is very low [26] and so it was not surprising that protein levels were below the limit of detection, given the small amount (1 µg) of total protein analysed. In any event there was no evidence of an extended spectrum SHV variant [26], and so the presence/absence of SHV did not influence the β-lactam susceptibility predictions made. The second, more serious issue was that in around 25% of isolates, a small amount of KPC was detected by LC-MS/MS, though only by matching a small number of peptides in each case. The only true KPC positive sample, as validated by WGS, matched to 26 peptides. Because of this disparity, which likely arises because there is some protein in these samples having a few peptides with significant identity to peptides from KPC, WGS data were used to defined KPC status. In all other respects the binary identification of β-lactamase carriage/non-carriage by LC-MS/MS matched the WGS data. Of course, the information richness of the LC-MS/MS data allows consideration of protein abundance as well as presence/absence, as will be discussed below.

First, we simply used the binary β-lactamase identification to predict cephalosporin and carbapenem susceptibility in the 40 *K. pneumoniae* isolates. Errors between predicted and determined susceptibility are marked in Table 3, the disc susceptibility data are presented in Table 4. Critical errors, where we predicted susceptibility but the isolate was non-susceptible, were seen for at least one β-lactam in 4/40 isolates. For isolates KP11, KP21 and KP30, the only error was wrongly predicting cefoxitin susceptibility. For KP13, errors were for cefoxitin, doripenem and ertapenem. LC-MS/MS has already shown that each of these four isolates overproduce the AcrAB-TolC efflux pump because of a loss of function mutation in *ramR* [27]. KP11 and KP13 both carry CTX-M, and recently we have reported that a *ramR* loss of function mutation enhances the ability of CTX-M to influence cefoxitin and some carbapenem susceptibility, presumably because of AcrAB-TolC overproduction [16]. Therefore, this explains the incorrect prediction of cefoxitin susceptibility in KP11 (which carries a carbapenemase, masking the ability of AcrAB-TolC hyperproduction to enhance CTX-M mediated reduced carbapenem susceptibility), and of cefoxitin, doripenem and ertapenem susceptibility in KP13. Isolates KP21 and KP30 were also incorrectly predicted to be cefoxitin susceptible. In these cases, they do not carry CTX-M, but instead carry TEM, so we conclude from this that TEM is also empowered to confer cefoxitin non-susceptibility in the presence of AcrAB-TolC hyperproduction, which has not previously been reported. The only other two isolates in this collection carrying *ramR* mutations, and hyperproducing AcrAB-TolC [27] are KP4, which is pan-β-lactam resistant because of CTX-M, OXA-232 and NDM-1 β-lactamases, so the effect of *ramR* mutation is not apparent, and KP59, which retains susceptibility to all test β-lactams, including cefoxitin. However, this isolate does not carry TEM or CTX-M, and we have recently shown that in the absence of a β-lactamase, loss of *ramR* does not mediate β-lactam non-susceptibility [16].

**Table 3:**
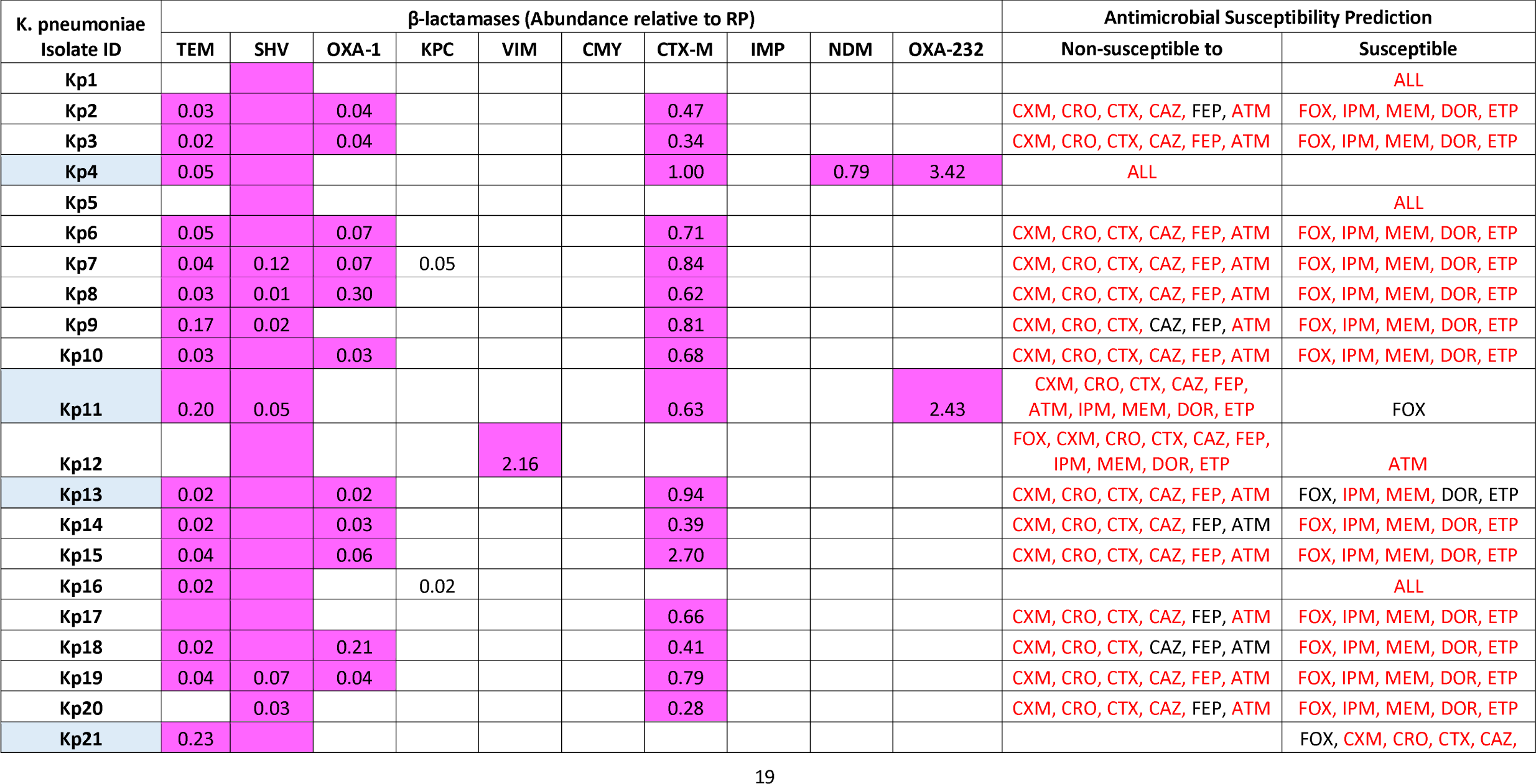
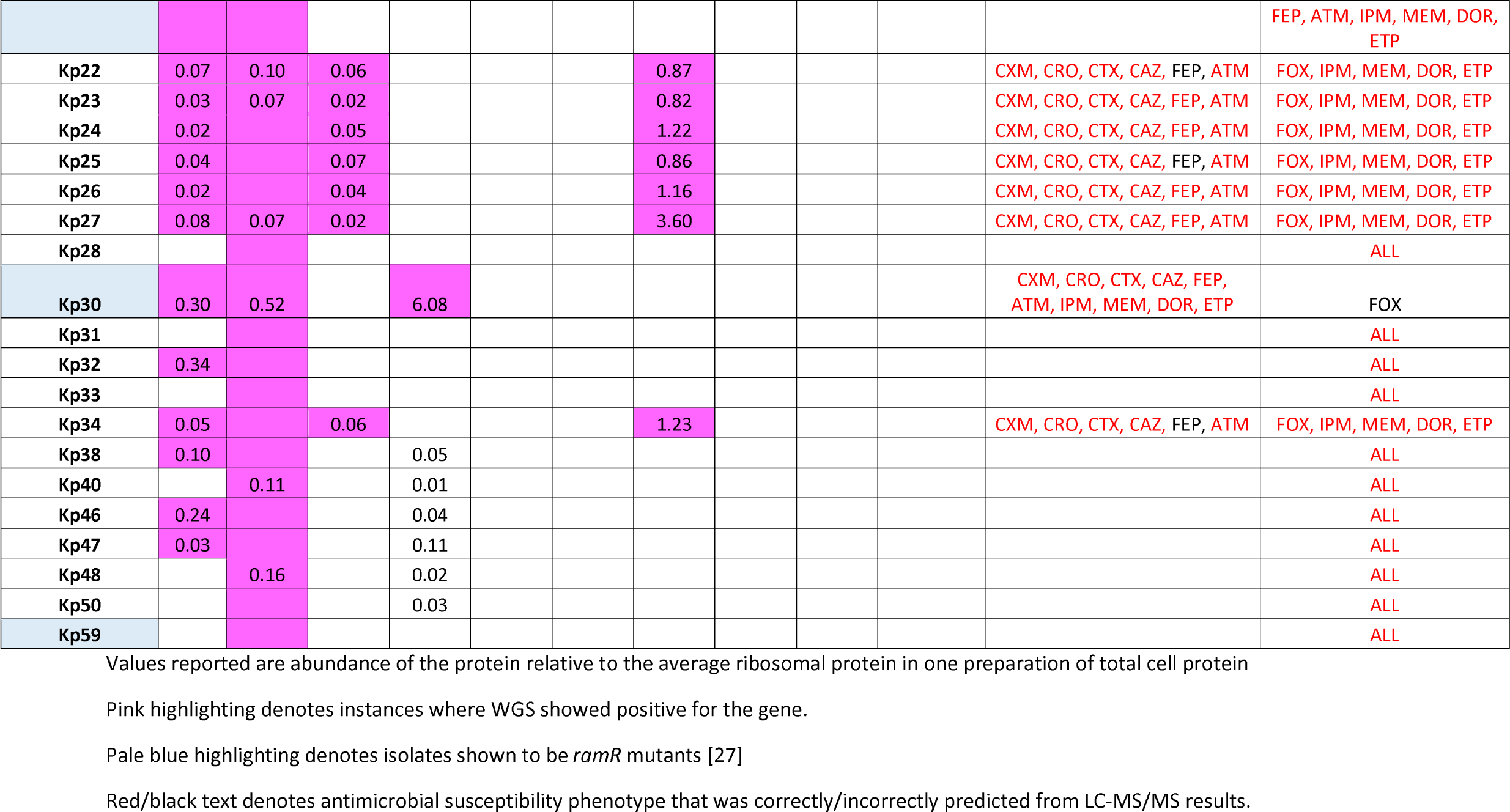
Prediction of β-lactam susceptibility in *K. pneumonia* clinical isolates

**Table 4.** Antimicrobial susceptibility testing for *K. pneumonia* clinical isolates.

| *K. pneumoniae*<br>Isolate ID | AST Results |  |  |  |  |  |  |  |  |  |  |
| --- | --- | --- | --- | --- | --- | --- | --- | --- | --- | --- | --- |
|  | FOX | CXM | CRO | CTX | CAZ | FEP | ATM | IPM | MEM | DOR | ETP |
| Kp1 | 26 | 21 | 32 | 32 | 29 | 32 | 33 | 26 | 28 | 27 | 31 |
| Kp2 | 25 | 6 | 12 | 11 | 20 | 21 | 18 | 25 | 29 | 25 | 28 |
| Kp3 | 24 | 6 | 10 | 10 | 16 | 17 | 10 | 26 | 29 | 27 | 28 |
| Kp4 | 6 | 6 | 6 | 6 | 6 | 6 | 6 | 10 | 6 | 6 | 6 |
| Kp5 | 18 | 18 | 29 | 28 | 28 | 30 | 30 | 27 | 29 | 27 | 30 |
| Kp6 | 25 | 6 | 9 | 11 | 18 | 17 | 15 | 26 | 28 | 26 | 29 |
| Kp7 | 23 | 6 | 10 | 11 | 18 | 16 | 14 | 24 | 28 | 24 | 28 |
| Kp8 | 25 | 6 | 9 | 8 | 17 | 15 | 13 | 24 | 30 | 24 | 30 |
| Kp9 | 26 | 6 | 11 | 11 | 21 | 19 | 19 | 27 | 29 | 25 | 30 |
| Kp10 | 25 | 6 | 12 | 13 | 20 | 18 | 17 | 24 | 29 | 27 | 30 |
| Kp11 | 7 | 6 | 6 | 6 | 14 | 9 | 8 | 16 | 10 | 11 | 6 |
| Kp12 | 6 | 6 | 8 | 6 | 6 | 6 | 27 | 11 | 8 | 8 | 6 |
| Kp13 | 8 | 6 | 6 | 6 | 11 | 6 | 6 | 25 | 23 | 21 | 17 |
| Kp14 | 26 | 6 | 11 | 13 | 20 | 21 | 20 | 25 | 30 | 25 | 30 |
| Kp15 | 23 | 6 | 8 | 9 | 16 | 13 | 14 | 25 | 29 | 24 | 27 |
| Kp16 | 26 | 23 | 30 | 31 | 28 | 31 | 33 | 26 | 29 | 27 | 30 |
| Kp17 | 24 | 6 | 10 | 10 | 19 | 19 | 17 | 27 | 30 | 27 | 29 |
| Kp18 | 25 | 6 | 12 | 14 | 23 | 21 | 22 | 25 | 30 | 27 | 27 |
| Kp19 | 25 | 6 | 10 | 10 | 18 | 18 | 17 | 25 | 29 | 26 | 27 |
| Kp20 | 22 | 6 | 14 | 16 | 20 | 22 | 17 | 25 | 29 | 26 | 28 |
| Kp21 | 16 | 18 | 27 | 29 | 26 | 29 | 29 | 28 | 31 | 28 | 28 |
| Kp22 | 25 | 6 | 10 | 12 | 18 | 19 | 17 | 25 | 29 | 26 | 28 |
| Kp23 | 26 | 6 | 11 | 11 | 18 | 17 | 17 | 25 | 29 | 26 | 28 |
| Kp24 | 25 | 6 | 8 | 10 | 18 | 17 | 14 | 25 | 30 | 26 | 27 |
| Kp25 | 27 | 6 | 10 | 12 | 19 | 21 | 18 | 26 | 30 | 26 | 27 |
| Kp26 | 24 | 6 | 8 | 9 | 17 | 16 | 12 | 25 | 30 | 25 | 27 |
| Kp27 | 24 | 6 | 6 | 6 | 6 | 8 | 6 | 28 | 30 | 28 | 24 |
| Kp28 | 21 | 20 | 29 | 30 | 27 | 31 | 32 | 29 | 30 | 28 | 29 |
| Kp30 | 6 | 6 | 8 | 10 | 6 | 7 | 6 | 10 | 6 | 6 | 6 |
| Kp31 | 23 | 21 | 30 | 27 | 29 | 31 | 31 | 25 | 25 | 25 | 25 |
| Kp32 | 23 | 22 | 30 | 28 | 29 | 31 | 32 | 27 | 24 | 27 | 25 |
| Kp33 | 24 | 23 | 30 | 32 | 30 | 32 | 33 | 27 | 30 | 27 | 29 |
| Kp34 | 23 | 6 | 8 | 10 | 15 | 19 | 13 | 26 | 27 | 27 | 23 |
| Kp38 | 20 | 21 | 28 | 28 | 28 | 30 | 31 | 27 | 30 | 28 | 30 |
| Kp40 | 23 | 24 | 30 | 30 | 28 | 31 | 34 | 26 | 29 | 27 | 29 |
| Kp46 | 23 | 22 | 29 | 31 | 26 | 30 | 33 | 26 | 31 | 28 | 29 |
| Kp47 | 26 | 25 | 31 | 31 | 28 | 31 | 32 | 25 | 30 | 26 | 28 |
| Kp48 | 24 | 23 | 29 | 31 | 27 | 30 | 32 | 25 | 30 | 27 | 28 |
| Kp50 | 25 | 27 | 33 | 33 | 29 | 31 | 33 | 25 | 30 | 25 | 30 |
| Kp59 | 28 | 21 | 30 | 32 | 28 | 30 | 32 | 26 | 33 | 28 | 31 |

Zone diameters are reported in mm, following standard CLSI methodology. Shading: Blue, susceptible, Green, non-susceptible (intermediate resistant), Yellow, resistant.

In terms of non-critical errors, these were seen commonly for cefepime (9/40), and less commonly for ceftazidime (2/40) and aztreonam (2/40), where in each case, non-susceptibility was incorrectly predicted in CTX-M producers (Table 3). WGS confirmed that each of these CTX-M enzymes was CTX-M15, so we presumed that there must be some background difference to explain the weaker effect of CTX-M15 carriage in these nine isolates. We considered OmpK35 and OmpK36 porin abundance and AcrAB efflux pump levels (which can all affect envelope permeability), and also the abundance of CTX-M itself. All abundance data for these proteins were extracted from the LC-MS/MS data and a pairwise comparison between the 9/21 cefepime susceptible versus the 12/21 cefepime resistant, CTX-M positive, carbapenemase negative isolates showed no significant differences in abundance between the groups. The data for CTX-M (cefepime resistant isolates have, on average 1.8 fold more CTX-M than cefepime susceptible isolates, p=0.06) and OmpK35 (cefepime susceptible isolates have, on average 1.6 fold less OmpK35 than cefepime susceptible isolates, p=0.09) were more promising than AcrAB (p=0.21) and OmpK36 (p=0.24). We found that if we crudely factored together CTX-M abundance with one of the permeability factors, either by subtracting porin abundance from the CTX-M abundance or by adding the efflux abundance to the CTX-M abundance, we observed that the combined influence of CTX-M upregulation plus OmpK35 downregulation was significantly implicated in cefepime resistance (p=0.02); the other permeability factors did not generate a significant effect (CTX-M/OmpK36, p=0.09; CTX-M/AcrAB, p=0.07). An association between CTX-M production and cefepime susceptibility has previously been made [28], but the additional impact of OmpK35 downregulation has not been noted before.

### Prediction of β-lactam susceptibility in E. coli, P. aeruginosa and A. baumannii

Given our success in integrating binary β-lactamase identification with abundance measurements for β-lactamases and proteins involved in envelope permeability to predict β-lactam susceptibility in *K. pneumoniae* without any critical errors, next we considered a small sample of three additional species (Table 5). As with *K. pneumoniae*, there were a small number of low abundance, low peptide hit calls for β-lactamases that were not supported by the WGS data in 3/15 isolates (Table 5). This suggests that we can apply even stricter cut-offs to our analysis. Only β-lactamase presence supported by WGS was used to predict antibacterial drug susceptibility. For *E. coli*, four isolates producing various β-lactamases, plus one pan-susceptible isolate, were considered. Isolate IR24, was flagged by the LC-MS/MS as carrying an extended spectrum β-lactamase (ESBL) SHV variant and this was confirmed by WGS to be SHV-12. Since we had not tested an SHV in our transformants (Table 2) we used the CTX-M *E. coli* transformant (another ESBL) to predict susceptibility in the SHV carrying isolate IR24. Because of this, we erroneously predicted ceftazidime susceptibility because SHV-12 is known to confer ceftazidime resistance [26]. All other predictions for *E. coli* were correct, with carbapenem resistance being seen in two isolates because of NDM-1 carriage (Table 5, Table 6).

**Table 5:** Prediction of β-lactam susceptibility in non-*K. pneumoniae* clinical isolates

| *E. coli* Isolate ID | β-lactamases (Abundance relative to RP) | | | | | | | | | Others (Porins, efflux, AmpC) | | | | | Antimicrobial Resistance Prediction | |
| --- | --- | --- | --- | --- | --- | --- | --- | --- | --- | --- | --- | --- | --- | --- | --- | --- |
|  | TEM | SHV | KPC | VIM | CMY | CTX-M | IMP | NDM | OXA-1 | OmpF | OmpC | AcrA | AcrB | TolC | Non-susceptible | Susceptible |
| ATCC 25922 |  |  |  |  |  |  |  |  |  | 2.49 | 2.21 | 0.17 | 0.08 | 0.18 |  | ALL |
| IR10 | 0.39 | 0.07 |  |  | 0.51 | 2.11 |  | 0.73 |  | 0.01 | 1.59 | 0.22 | 0.08 | 0.08 | ALL |  |
| IR15 |  | 0.17 |  |  |  | 1.89 |  |  | 0.13 |  | 0.53 | 0.07 | 0.02 | 0.10 | CXM, CRO, CTX, CAZ, FEP, ATM | FOX, IPM, MEM, DOR, ETP |
| IR24 |  | 0.72 |  |  |  |  |  |  |  | 0.92 | 0.71 | 0.08 | 0.03 | 0.12 | CXM, CRO, CTX, FEP, ATM | FOX, CAZ, IPM, MEM, DOR, ETP |
| IR60 | 0.10 | 0.10 | 0.10 |  |  | 0.75 | 0.08 | 0.28 | 0.01 | 0.51 | 1.57 | 0.11 | 0.08 | 0.11 | ALL |  |

| *P. aeruginosa* Isolate ID | β-lactamases (Abundance relative to RP) | | | | | | | | | Others (Porins, efflux, AmpC) | | | | | Antimicrobial Resistance Prediction | |
| --- | --- | --- | --- | --- | --- | --- | --- | --- | --- | --- | --- | --- | --- | --- | --- | --- |
|  | TEM | SHV | KPC | VIM | CMY | CTX-M | IMP | NDM | OXA-2 | OprD | MexA | MexB | OprM | AmpC | Non-susceptible | Susceptible |
| 81-11963 |  |  |  | 0.46 |  |  |  |  |  |  | 0.16 | 0.06 | 0.09 | 0.07 | CAZ, FEP, IPM, MEM, DOR | ATM |
| 404-00 |  |  |  | 0.71 |  |  |  |  |  | 0.19 | 0.15 | 0.05 | 0.11 |  | CAZ, FEP, IPM, MEM, DOR | ATM |
| 301-5473 |  |  |  | 0.22 |  |  |  |  | 0.27 |  | 0.61 | 0.15 | 0.33 | 0.06 | CAZ, FEP, IPM, MEM, DOR | ATM |
| 86-14571 |  |  |  |  |  |  | 0.13 |  |  |  | 1.56 | 0.47 | 0.76 | 16.19 |  | CAZ, FEP, ATM, IPM, MEM, DOR |
| 73-56826 |  |  | 0.02 |  |  |  |  |  |  | 0.17 | 1.02 | 0.34 | 0.45 | 10.67 |  | CAZ, FEP, ATM, IPM, MEM, DOR |

| *A. baumannii* Isolate ID | β-lactamases (Abundance relative to RP) | | | | | | | | | Others (Efflux, OXA-51) | | | | | Antimicrobial Resistance Prediction | |
| --- | --- | --- | --- | --- | --- | --- | --- | --- | --- | --- | --- | --- | --- | --- | --- | --- |
|  | TEM | SHV | KPC | VIM | CMY | CTX-M | IMP | NDM | OXA-1 | OXA-51 | AdeA | AdeB | AdeJ | AdeK | Non-susceptible | Susceptible |
| 4034C | 0.24 |  |  |  |  |  |  |  |  | 3.31 | 0.31 | 0.01 | 0.11 | 0.27 |  | ALL |
| 4737C | 0.02 |  |  |  |  |  |  |  |  | 4.66 | 0.36 | 0.04 | 0.12 | 0.32 |  | ALL |
| **4742C** | 0.01 |  |  |  |  |  |  |  |  | 1.95 | 0.30 | 0.10 | 0.11 | 0.26 |  | ALL |
| **4750C** | 0.01 |  |  |  |  |  |  |  |  | 1.84 | 0.28 | 0.03 | 0.09 | 0.28 |  | ALL |
| **6490A** | 0.30 | 0.08 |  |  |  |  |  |  |  | 3.88 | 0.30 | 0.01 | 0.14 | 0.30 |  | ALL |

Values reported are abundance of the protein relative to the average ribosomal protein in one preparation of total cell protein
Pink highlighting denotes instances where WGS showed positive for the gene.
Pale blue highlighting denotes isolates shown to be *ramR* mutants [27]
Red/black text denotes antimicrobial susceptibility phenotype that was correctly/incorrectly predicted from LC-MS/MS results.

**Table 6.** Antimicrobial susceptibility testing for non-*K. pneumoniae* clinical isolates.

| *E. coli* Isolate ID | AST Results |  |  |  |  |  |  |  |  |  |  |
| --- | --- | --- | --- | --- | --- | --- | --- | --- | --- | --- | --- |
|  | FOX | CXM | CRO | CTX | CAZ | FEP | ATM | IPM | MEM | DOR | ETP |
| ATCC 25922 | 30 | 25 | 33 | 33 | 30 | 32 | 33 | 25 | 29 | 29 | 37 |
| IR10 | 6 | 6 | 6 | 6 | 6 | 6 | 6 | 9 | 6 | 6 | 6 |
| IR15 | 25 | 6 | 6 | 6 | 9 | 15 | 6 | 27 | 28 | 26 | 26 |
| IR24 | 25 | 15 | 17 | 17 | 11 | 23 | 10 | 26 | 27 | 23 | 24 |
| IR60 | 6 | 6 | 6 | 6 | 6 | 12 | 9 | 18 | 17 | 16 | 16 |

| *P. aeruginosa* Isolate ID | AST Results |  |  |  |  |  |  |  |  |  |  |
| --- | --- | --- | --- | --- | --- | --- | --- | --- | --- | --- | --- |
|  | FOX | CXM | CRO | CTX | CAZ | FEP | ATM | IPM | MEM | DOR | ETP |
| 81-11963 | 6 | 6 | 6 | 6 | 10 | 13 | 22 | 6 | 6 | 6 | 6 |
| 404-00 | 6 | 6 | 6 | 6 | 14 | 12 | 28 | 8 | 14 | 11 | 6 |
| 301-5473 | 6 | 6 | 6 | 6 | 12 | 13 | 20 | 6 | 6 | 6 | 6 |
| 86-14571 | 6 | 6 | 6 | 6 | 6 | 6 | 13 | 6 | 6 | 6 | 6 |
| 73-56826 | 6 | 6 | 6 | 6 | 10 | 15 | 8 | 24 | 25 | 25 | 9 |

| *A. baumannii* Isolate ID | AST Results |  |  |  |  |  |  |  |  |  |  |
| --- | --- | --- | --- | --- | --- | --- | --- | --- | --- | --- | --- |
|  | FOX | CXM | CRO | CTX | CAZ | FEP | ATM | IPM | MEM | DOR | ETP |
| 4034C | 6 | 6 | 6 | 6 | 6 | 17 | 8 | 11 | 13 | 15 | 6 |
| 4737C | 6 | 6 | 6 | 6 | 6 | 14 | 12 | 9 | 13 | 14 | 6 |
| 4742C | 6 | 6 | 6 | 6 | 10 | 12 | 6 | 9 | 14 | 16 | 6 |
| 4750C | 6 | 6 | 6 | 6 | 6 | 13 | 9 | 10 | 14 | 16 | 6 |
| 6490A | 6 | 6 | 6 | 6 | 6 | 15 | 6 | 10 | 12 | 15 | 6 |

Zone diameters are reported in mm, following standard CLSI methodology. Shading: Blue, susceptible, Green, non-susceptible (intermediate resistant), Yellow, resistant. Grey – assumed to be intrinsically resistant, so no breakpoint defined.

In *P. aeruginosa*, VIM and IMP production was seen in 3/5 and 1/5 isolates respectively (Table 5), meaning that aztreonam susceptibility was predicted, since VIM and IMP do not confer aztreonam resistance (Table 2) and there was no other mobile aztreonam hydrolysing β-lactamase present. This prediction of aztreonam susceptibility proved incorrect in isolates 301-5473 (VIM producer) and 86-14571 (IMP producer) (Table 6). According to the LC-MS/MS data, these isolates hyperproduce the MexAB-OprM efflux pump, relative to PA01, and isolates 81-11963 and 404-00, which also carry VIM, and yet are aztreonam susceptible, as predicted (Table 5). We conclude, therefore that MexAB-OprM hyperproduction is at least partly responsible for aztreonam non-susceptibility in isolates 301-5473 and 86-14571, though the latter also hyper-produces its chromosomal AmpC β-lactamase, intrinsic to *P. aeruginosa* and also able to hydrolyse aztreonam (Table 5). Isolate 73-56826 does not carry any acquired β-lactamases, so, based on the susceptibility profile of PA01, it was predicted to be susceptible to all β-lactams (Table 2), but in fact it is non-susceptible to ceftazidime, cefepime and aztreonam (Table 6). Again, the LC-MS/MS data allowed us to identify that this isolate hyper-produces the chromosomal AmpC β-lactamase and MexAB-OprM, again explaining cephalosporin resistance. LC-MS/MS revealed that several of the carbapenem resistant isolates produce no detectable OprD porin, which is produced at wild-type levels in the carbapenem susceptible, AmpC hyperproducing isolate 73-56826 (Table 5). The involvement of AmpC hyperproduction, MexAB-OprM hyperproduction and OprD downregulation in cephalosporin and carbapenem resistance in *P. aeruiginosa* clinical isolates are all well-known phenomena [29], but each is under complex control [30], and so currently it is not possible to predict abundance for these proteins based solely on WGS [14] and *P. aeruginosa* is considered very challenging even for proteomics based antimicrobial susceptibility prediction [17]. Our LC-MS/MS methodology shows that it is possible to achieve.

In the A. *baumannii* isolates, there was no evidence of any acquired β-lactamase genes other than the ampicillin hydrolysing TEM enzyme in the LC-MS/MS data, and this was confirmed by WGS (Table 5). However, all isolates were found to be non-susceptible to all test β-lactams, including the carbapenems (Table 6). Analysis of the LC-MS/MS data again identified the reason, confirming that all isolates hyperproduce their chromosomally encoded, intrinsic OXA-51-like carbapenemase, relative to the control isolate (Table 1), in which OXA-51 production was below the level of detection in the LC-MS/MS. OXA-51 is known to confer pan-β-lactam resistance when hyperproduced in A. *baumannii* [31].

## Conclusions

We have, for the first time, reported a comprehensive analysis of the ability of LC-MS/MS to be used as a tool to collect protein identification and abundance data for the prediction of ABR in multi-drug resistant bacterial isolates. We have shown that our methodology is robust and can successfully predict antimicrobial susceptibility using a single run. Even when using only 1 µg of total cell protein, and applying a very strict cut-off, that only proteins represented by three or more peptides were included, approximately 1000 proteins were identified and quantified in each sample, and clearly this means that we may miss some detail. However, the key ABR determinants are expressed at high abundance, and WGS comparisons actually suggest that we could increase the strictness of our cut-off peptide numbers to reduce the number of false positive hits seen for some β-lactamases in the LC-MS/MS data from some samples. Currently, our analysis requires cultured bacteria. Once a culture has been grown, the process of extraction, SDS-PAGE, tryptic digestion, LC-MS/MS and data analysis takes around 24 hours. This is not competitive even with existing culture based antimicrobial susceptibly testing. However, it is possible that proteins extracted directly from clinical samples – e.g. from culture positive blood via the Bruker Sepsityper system [32] – can be analysed directly via LC-MS/MS. This would reduce time to susceptibility testing by around 12 hours from the current situation but such is the severity of bloodstream infection that even this relatively modest improvement in time to susceptibility testing result might have significant impact on patient wellbeing [32]. It must be accepted that our methodology does not lend itself to being run as a high throughput system, so perhaps these rare and serious infections are the most reasonable place to deploy it. On the other hand, it may be that the future for this methodology is to improve our understanding of how antibacterial susceptibility phenotype is influenced by bacterial genotype, bridging the gap in our understanding of which SNPs cause which proteins to be differentially produced, and which levels of production are necessary to alter antibacterial drug susceptibility. Since WGS methodologies have the potential to be more widely used direct from clinical samples, in a higher throughput way, if biological interpretation of WGS data could be made more complete, the potential for revolutionising patient care would be particularly great [14].

## References

[1] DOH/DEFRA (2013) UK 5 Year Antimicrobial Resistance Strategy 2013-2018. www.gov.uk/government/publications

[2] Rice LB. (2010) Infect Control Hosp Epidemiol. 31 Suppl 1:S7–10.

[3] Pea F, Viale P (2009) Crit Care. 13:214.

[4] Bassetti M, Righi E. (2013) Hematology Am Soc Hematol Educ Program. 2013:428–32.

[5] Karam G, Chastre J, Wilcox MH, Vincent JL. Antibiotic strategies in the era of multidrug resistance. Crit Care. 2016 Jun 22;20(1):136.

[6] Didelot X, Bowden R, Wilson DJ et al. Transforming clinical microbiology with bacterial genome sequencing. Nat Rev Genet 2012; 13: 601–12.

[7] Livermore DM, Wain J. Revolutionising bacteriology to improve treatment outcomes and antibiotic stewardship. Infect Chemother 2013; 45: 1–10.

[8] Gordon NC, Price JR, Cole K et al. Prediction of Staphylococcus aureus antimicrobial resistance by whole-genome sequencing. J Clin Microbiol 2014; 52: 1182–91.

[9] Wilson MR, Naccache SN, Samayoa E et al. Actionable diagnosis of neuroleptospirosis by next-generation sequencing. N Engl J Med 2014; 370: 2408–17.

[10] Grumaz S, Stevens P, Grumaz C et al. Next-generation sequencing diagnostics of bacteremia in septic patients. Genome Med 2016; 8: 73.

[11] Hasman H, Saputra D, Sicheritz-Ponten T et al. Rapid whole-genome sequencing for detection and characterization of microorganisms directly from clinical samples. J Clin Microbiol 2014; 52: 139–46.

[12] Schmidt K, Mwaigwisya S, Crossman LC, Doumith M, Munroe D, Pires C, Khan AM, Woodford N, Saunders NJ, Wain J, O’Grady J, Livermore DM. Identification of bacterial pathogens and antimicrobial resistance directly from clinical urines by nanopore-based metagenomic sequencing. J Antimicrob Chemother. 2017 Jan;72(1):104–114.

[13] Stoesser N, Batty EM, Eyre DW, Morgan M, Wyllie DH, Del Ojo Elias C, et al. Predicting antimicrobial susceptibilities for Escherichia coli and Klebsiella pneumoniae isolates using whole genomic sequence data J Antimicrob Chemother, 68 (2013), pp. 2234–2244

[14] Ellington MJ, Ekelund O, Aarestrup FM, Canton R, Doumith M, Giske C, Grundman H, Hasman H, Holden MT, Hopkins KL, Iredell J, Kahlmeter G, Köser CU, MacGowan A, Mevius D, Mulvey M, Naas T, Peto T, Rolain JM, Samuelsen Ø, Woodford N. The role of whole genome sequencing in antimicrobial susceptibility testing of bacteria: report from the EUCAST subcommittee. Clin Microbiol Infect. 2017 Jan;23(1):2–22.

[15] Alekshun MN, Levy SB. (2007). Cell. 128:1037–50

[16] Jimenez-Castellanos, JC, Wan Nur Ismah WAK, Takebayashi, Y, Findlay J, Schneiders T, Heesom, KJ, Avison, MB. The Envelope Proteome Changes Driven By RamA Overproduction in Klebsiella pneumoniae That Enhance Acquired β-Lactam Resistance BioRxiv 2017. doi: https://doi.org/10.1101/133918

[17] Charretier Y, Schrenzel J. Mass spectrometry methods for predicting antibiotic resistance. Proteomics Clin Appl. 2016 Oct;10(9–10):964–981.

[18] Jiménez-Castellanos JC, Wan Ahmad Kamil WN, Cheung CH, Tobin MS, Brown J, Isaac SG, Heesom KJ, Schneiders T, Avison MB. Comparative effects of overproducing the AraC-type transcriptional regulators MarA, SoxS, RarA and RamA on antimicrobial drug susceptibility in Klebsiella pneumoniae. J Antimicrob Chemother. 2016 Jul;71(7):1820–5.

[19] Clinical and Laboratory Standards Institute. Performance Standards for Antimicrobial Disk Susceptibility Tests – Ninth Edition: Approved Standard M2-A9. CLSI, Wayne, PA, USA. 2006.

[20] Clinical and Laboratory Standards Institute. Performance Standards for Antimicrobial Susceptibility Testing: Twenty-fourth Informational Supplement M100-S24. CLSI, Wayne, PA, USA. 2014.

[21] Kuo SC, Yang SP, Lee YT, Chuang HC, Chen CP, Chang CL, Chen TL, Lu PL, Hsueh PR, Fung CP. Dissemination of imipenem-resistant Acinetobacter baumannii with new plasmid-borne blaOXA-72 in Taiwan. BMC Infect Dis. 2013 Jul 13; 13: 319

[22] Silva JC, Gorenstein M V, Li G-Z, Vissers JPC, Geromanos SJ. Absolute quantification of proteins by LCMSE: a virtue of parallel MS acquisition. Mol Cell Proteomics. 2006;5(1):144–156.

[23] Bolger AM, Lohse M Usadel B. Trimmomatic: a flexible trimmer for Illumina sequence data. Bioinformatics 2014; 30: 2114–20.

[24] Zankari E, Hasman H, Cosentino S et al. Identification of acquired antimicrobial resistance genes. J Antimicrob Chemother 2012; 67: 2640–4.

[25] Carattoli A, Zankari E, Garcia-Fernandez A et al. In silico detection and typing of plasmids using PlasmidFinder and plasmid multilocus sequence typing. Antimicrob Agents Chemother 2014; 58: 3895–903.

[26] Ford PJ, Avison MB. Evolutionary mapping of the SHV beta-lactamase and evidence for two separate IS26-dependent blaSHV mobilization events from the Klebsiella pneumoniae chromosome. J Antimicrob Chemother. 2004 Jul;54(1):69–75.

[27] Wan Nur Ismah WAK,1,2 Takebayashi Y, Findlay J, Heesom KJ, Jiménez-Castellanos JC, Zhang J, Graham L, Bowker K, Williams OM, MacGowan AP and Avison MB. Prediction of fluoroquinolone susceptibility directly from whole genome sequence data using liquid chromatography-tandem mass spectrometry to identify mutant genotypes. bioRxiv 2017.

[28] Ma L, Siu LK, Lu PL. Effect of spacer sequences between bla(CTX-M) and ISEcp1 on bla(CTX-M) expression. J Med Microbiol. 2011 Dec;60(Pt 12):1787–92.

[29] Castanheira M, Mills JC, Farrell DJ, Jones RN. Mutation-driven β-lactam resistance mechanisms among contemporary ceftazidime-nonsusceptible Pseudomonas aeruginosa isolates from U.S. hospitals. Antimicrob Agents Chemother. 2014 Nov;58(11):6844–50.

[30] Lister PD, Wolter DJ, Hanson ND. Antibacterial-resistant Pseudomonas aeruginosa: clinical impact and complex regulation of chromosomally encoded resistance mechanisms. Clin Microbiol Rev. 2009 Oct;22(4):582–610.

[31] Evans BA, Amyes SG. OXA β-lactamases. Clin Microbiol Rev. 2014 Apr;27(2):241–63.

[32] Morgenthaler NG, Kostrzewa M. Rapid identification of pathogens in positive blood culture of patients with sepsis: review and meta-analysis of the performance of the sepsityper kit. Int J Microbiol. 2015;2015:827416.

